# Sensory origin of oculomotor variability revealed by MT population activity

**DOI:** 10.64898/2026.07.14.738423

**Authors:** Hoi Ming Ken Yip, Shaun Liam Cloherty, Maureen Ann Hagan, Nicholas Seow Chiang Price

## Abstract

Motor outputs vary even in response to identical sensory inputs, yet the origin of this variability within the sensorimotor pathway remains unresolved. Here, we evaluate how variability in sensory representations can explain behavioural fluctuations in a reflexive oculomotor task. We used Neuropixels probes to record neuronal activity in area MT of marmosets during ocular following responses and applied partial least squares regression to extract the shared variance between population activity and eye movements. Stimulus-evoked activity reliably predicted trial-by-trial variability in open-loop eye velocity. Additionally, closed-loop analyses revealed trial-by-trial correspondence between eye movements and subsequent neural responses. These results demonstrate that variability in sensory neural populations contributes to motor variability, supporting the claim that sensory noise is propagated through the sensorimotor pathway. Surprisingly, however, traditional models trained to only capture neural variability across trials poorly predicted behavioural variations, suggesting that only a subset of sensory representations is accessible to the motor system.

## Introduction

Motor behaviours are inherently variable. Even elite athletes, despite exceptional skill and experience, cannot reproduce identical actions across repetitions. This variability persists under tightly controlled laboratory conditions, where identical sensory inputs nonetheless yield variable perceptual and behavioural responses in both humans and animals (Afshar et al., 2011; Churchland et al., 2006; Osborne et al., 2005). While much behavioural variability must arise from internal fluctuations within the sensorimotor pathway, it is common to study this variability using tasks that require volitional behaviours, making it difficult to disambiguate the influence of top-down influences such as motivation and attention. A central question, therefore, is to determine what accounts for variability in time-critical, reflexive behaviours that depend on complex neural processing.

One hypothesis attributes behavioural variability to the stochastic nature of neural firing (Faisal et al., 2008), proposing that variability is progressively averaged out and reduced through successive stages of processing (Maimon & Assad, 2009). Accordingly, neural variability should exert a greater influence on behaviour if it is present in downstream motor areas than in early sensory regions (Churchland et al., 2006; van Beers et al., 2004). Alternatively, neural variability evident at the level of sensory encoding may persist through the sensorimotor hierarchy in a form that neither grows nor diminishes; therefore it appears that noise in downstream areas exerts comparatively little additional influence on behavioural variability (Osborne et al., 2005). The key distinction between these views lies in how strongly variability in sensory representations correlates with behavioural outcomes.

The oculomotor system provides a powerful model for addressing this issue due to its continuous behavioural readouts and well-characterized neural circuitry. Previous single-neuron studies have revealed trial-by-trial correlations between smooth pursuit eye movements and firing rates in the middle temporal area (MT), a motion-sensitive visual region (Hohl et al., 2013; Lee et al., 2016). Notably, MT activity up to 60 ms before pursuit onset predicts subsequent movement kinematics, suggesting that sensory encoding variability can influence motor output. However, single-neuron analyses may overlook properties that only become evident at the population level (Ebitz & Hayden, 2021; Kohn et al., 2016). For instance, while single neurons in the motor cortex do not exhibit consistent tuning to motion parameters (Churchland et al., 2010; Churchland & Shenoy, 2007), motor response and planning can be predicted with population-level dynamics (Shenoy et al., 2013).

Here, we studied marmosets performing ocular following responses, in which the eyes reflexively track wide-field visual motion with extremely short latency (Miles et al., 1986; Pattadkal et al., 2023; Yip et al., 2023). Using a reflexive behaviour minimises top-down influences on sensory processing and tightly constrains the time windows of neural activity that can affect behaviour.

Using Neuropixels probes, we simultaneously recorded large neuronal populations in area MT and applied partial least squares (PLS) regression to predict instantaneous eye speed and direction on a single-trial basis. PLS, a reduced-rank regression technique, identifies latent components that jointly explain variability in neural activity and behaviour. Compared with other approaches, PLS offers several advantages: it mitigates multicollinearity; extracts behaviourally relevant components rather than dominant neural variance (as in principal component regression, PCR); and unlike hypothesis-driven methods such as vector averaging (Hohl et al., 2013; Huang & Lisberger, 2009), it incorporates the entire recorded population, including weakly tuned neurons.

We successfully modelled single-trial open-loop eye speeds with MT population activities before eye movement onset (mean r = 0.16-0.19). Moreover, PLS models computed with early eye speeds and later neural activities were also significant (mean r = 0.2-0.29), reflecting a closed-loop system in which population activity is affected by eye movement itself. Notably, PLS outperformed PCR and vector averaging, capturing neural variability more aligned with behavioural fluctuations despite explaining less total variance.

Together, these findings demonstrate that population activity in a sensory area incorporates trial-by-trial fluctuations that are predictive of behavioural variability in a reflexive oculomotor behaviour. By leveraging high-density recordings and population-level analyses, we reveal that sensory cortical variability can exert a measurable influence on downstream motor behaviour.

## Results

To investigate the relationship between sensory variability and motor variability, we recorded neural population activity in the middle temporal (MT) area during ocular following in two marmosets. Animals initiated a trial by fixating a peripheral target and subsequently made a saccade to a central target, which triggered the motion of a large-field background stimulus (Figure 1A). The sudden motion of the stimulus evoked an ocular following response. The direction of the background stimulus motion on each trial was chosen from 8 evenly-distributed directions. In 11/22 sessions (Monkey B) and 8/18 sessions (Monkey M) all directions were equally likely (all-directions task); in the remaining sessions, the stimulus direction of 0° was overrepresented, occurring with a 65% probability (biased-direction task). Data from the two tasks were pooled as there were no systematic differences in model performance between them. Neural activity in MT was recorded using a Neuropixels probe inserted to span all cortical layers. Neural data was spike-sorted with Kilosort 4 (Pachitariu et al., 2024). To retain as much information as possible from the whole population, we included both well-isolated neurons and multiunit activities in our analyses and considered both as “units” in the following sections.

**Figure 1.**
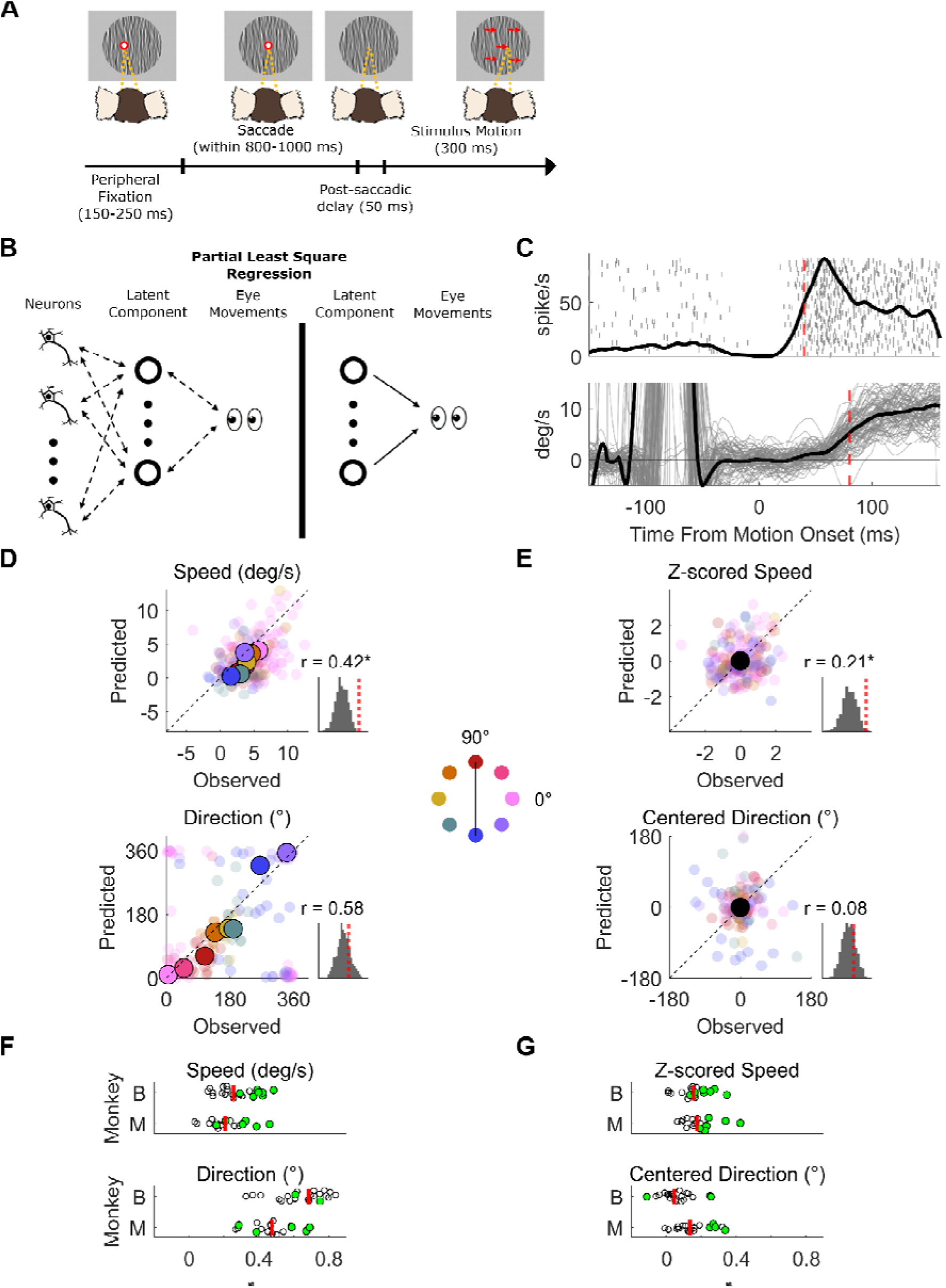
Open loop model to reconstruct eye speed and direction from middle temporal (MT) population activity before eye movement onset. (A) Experimental paradigm. Each trial began with the presentation of a stationary large field stimulus (diameter = 28°) and a peripheral fixation target located 5° from the center of the screen. Monkeys were required to fixate the peripheral target for 150-250 ms. Upon successful fixation, the target disappeared and reappeared centrally, prompting a saccadic movement. If the saccade was completed within 800-1000 ms, the fixation target disappeared, and the stimulus moved for 300 ms following a 50 ms delay, inducing ocular following responses. Stimuli were presented in eight directions, spaced 45° apart. (B) A schematic illustrating Partial Least Squares (PLS) regression. Latent components are extracted to explain variance in both neural activity and eye movements. In the regression step, these latent components are then used to reconstruct eye movements. (C) Upper panel: Raster plot of an example unit, with trial averaged peristimulus time histogram overlay. The red dotted line indicates the time point used in the model (40 ms). Lower panel: single trial eye speed (grey lines) across time in an example session, with trial averaged speed overlay (black thick line). The red dotted line indicates the time point used in the model (80 ms). (D) Scatter plots illustrating the relationship between observed and PLS reconstructed eye speed (upper panel) and direction (lower panel) at 80 ms, based on neural activity recorded at 40 ms in an example session. Each small dot represents an individual trial, with colors indicating different stimulus directions. Larger dots denote the mean values for each stimulus condition. The inset histograms show the distribution of correlation coefficients obtained from shuffled data, with the red dotted line indicating the experimental values. (E) Scatter plots of predicted and observed eye speed and direction after centering values within each stimulus condition to remove stimulus direction dependent effects. (F) Distribution of correlations in speed and direction in all recording sessions in two animals. Each dot represents one session (n_B_ = 22; n_M_ = 18). Green dots indicate significant correlation in that session. The red bar shows the median across sessions. (G) Similar to (F), but with z-scored speed and centered directions.

### Single-trial, open-loop eye movements successfully reconstructed with early neural activity

To reconstruct the speed and direction of eye movements using MT neural population activity, we employed Partial Least Squares (PLS) regression (Figure 1B). PLS was chosen for its ability to extract latent components that account for variance in both predictor and response variables (Abdi, 2010). The firing rates (convolved with Gaussian and discretized in 5-ms bins) of all simultaneously recorded neurons served as predictors, while horizontal and vertical eye velocities were used as response variables. Data from each recording session were analysed separately and we included trials from all stimulus conditions regardless of the number of trials within each condition. Using the PLS-predicted horizontal and vertical velocities (obtained with 10-fold cross validation), we computed eye speed and direction and evaluated the model performance based on the trial-by-trial correlation between these predictions and the observed values.

In our open-loop models, we reconstructed eye speed and direction at 80 ms after motion onset with neural activity recorded at 40 ms (Figure 1C). Since eye movements are initiated ∼50-60 ms after motion onset (Mean latency = 57.2 ms, SD = 4.76 ms), this analysis tests whether neural activities before eye movement initiation can predict eye speed and direction shortly after movement onset. This is the open loop phase of the visual response before the generated eye movements would start impacting neural responses. A previous study of smooth pursuit in macaques, in which similar predictions were made, focused on time windows of 20-60 and 80-120 ms post stimulus onset for spiking and eye speeds respectively (Hohl et al.,2013). The neural time window used in our analysis was centred around the same period used in that study, while our behavioural time windows were earlier. This adjustment reflects the shorter processing time due to differences in species and in model behaviour, eye movement onset is shorter on average in ocular following than smooth pursuit (Miles et al. 1986; Lee et al., 2016).

In a representative session in Monkey B, the reconstructed and observed eye speeds were significantly correlated (r = 0.42, p < 0.001, Figure 1C). Significance was assessed by comparison with a distribution of correlations generated by running PLS with shuffled trial labels within each stimulus direction 10,000 times. The correlation for eye direction, although high (r = 0.58), was not significantly different from the shuffled distribution (p = 0.24) (Figure 1D). Further analysis revealed that observed eye speed and direction differed significantly across stimulus conditions (Speed: one-way ANOVA, F(7,178) = 8.29, p < 0.001; Direction: Watson-Williams multi-sample test, F(7,178) =35.49, p < 0.001). This suggests that the high correlations between model predictions and observed data primarily reflects the model’s ability to distinguish between different stimulus directions rather than single trials within a condition.

To better account for behavioural biases and isolate trial-by-trial variability independent of stimulus direction, we removed stimulus direction dependent effects by centering single-trial responses relative to the mean of each condition. Specifically, for eye speed, we computed z-scores within each stimulus direction, while for eye direction, we calculated circular distance from the condition mean (Figure 1E). Following this normalization, the correlation between reconstructed and observed z-scored eye speeds remained significant for this session (r = 0.21, p = 0.001), while correlation for centered direction was also positive but not significant (r = 0.08, p = 0.22). These findings indicate that PLS captured trial-by-trial variations in eye speed irrespective of stimulus direction.

Across sessions, we were able to reconstruct eye speed and direction in both animals. For each of the four measures, the average model performance across all sessions (Fisher transformed mean correlation) was significant (Monkey B: t = 3.18-15.28, df = 21, all adjusted p < 0.01; Monkey M: t = 6.3-13.45, df = 17, all adjusted p < 0.01; corrected for multiple comparison). For eye speed (Figure 1E), 32% and 39% of sessions were significantly correlated in Monkey B and M, respectively. We found 9% and 28% of sessions had significant correlations for direction (Figure 1F). After removing stimulus direction dependent effects, 36% and 39% of sessions were significantly correlated for z-scored speed, and 14% and 22% were significant for centered direction (Figure 1G). We also examined model performance separately for sessions associated with the all-direction task and the biased-direction task. Supporting our decision to pool the two task types for analysis, there were no significant differences between correlations for the two tasks for any of the four measures (Monkey B: t = −1.14-1.45, df = 20, adjusted p = 0.65-0.84; Monkey M: t = −0.2-0.94, df = 16, adjusted p = 0.08-0.84, corrected for multiple comparison).

Closed-loop models suggest eye movements affect later neural activities on a trial-by-trial basis Oculomotor behaviour is a closed-loop system: when the eyes start tracking an object, it reduces the object’s speed on the retina. This altered visual input should subsequently affect neural responses in visual areas like MT. Unlike steady-state tracking in smooth pursuit where perfect tracking reduces the retinal speed of the tracked object to zero (Behling & Lisberger, 2020), eye speed in the ocular following response rarely reaches the stimulus speed before the subject makes a saccade. In fact, most studies on ocular following have focused only on the open-loop period, when gain is low (Kawano et al., 1994; Miles et al., 1986; Miura et al., 2014). To explore the effect of eye movements on later neural activities, we tested whether we could observe a trial-by-trial relationship between the two.

We approached the closed loop question by generating PLS models with early eye movements (80 ms) and later neural activities (120 ms). As before, we evaluated the model performance using the correlation between reconstructed and observed eye speeds and directions (Figure 2A, B). In the example session, we found significant correlation in eye speed (r = 0.54, p < 0.01) and z-scored speed (r = 0.31, p < 0.01). The correlation for direction and centered direction were also significant (direction: r = 0.76, p = 0.049, centered direction: r = 0.28, p = 0.045).

**Figure 2.**
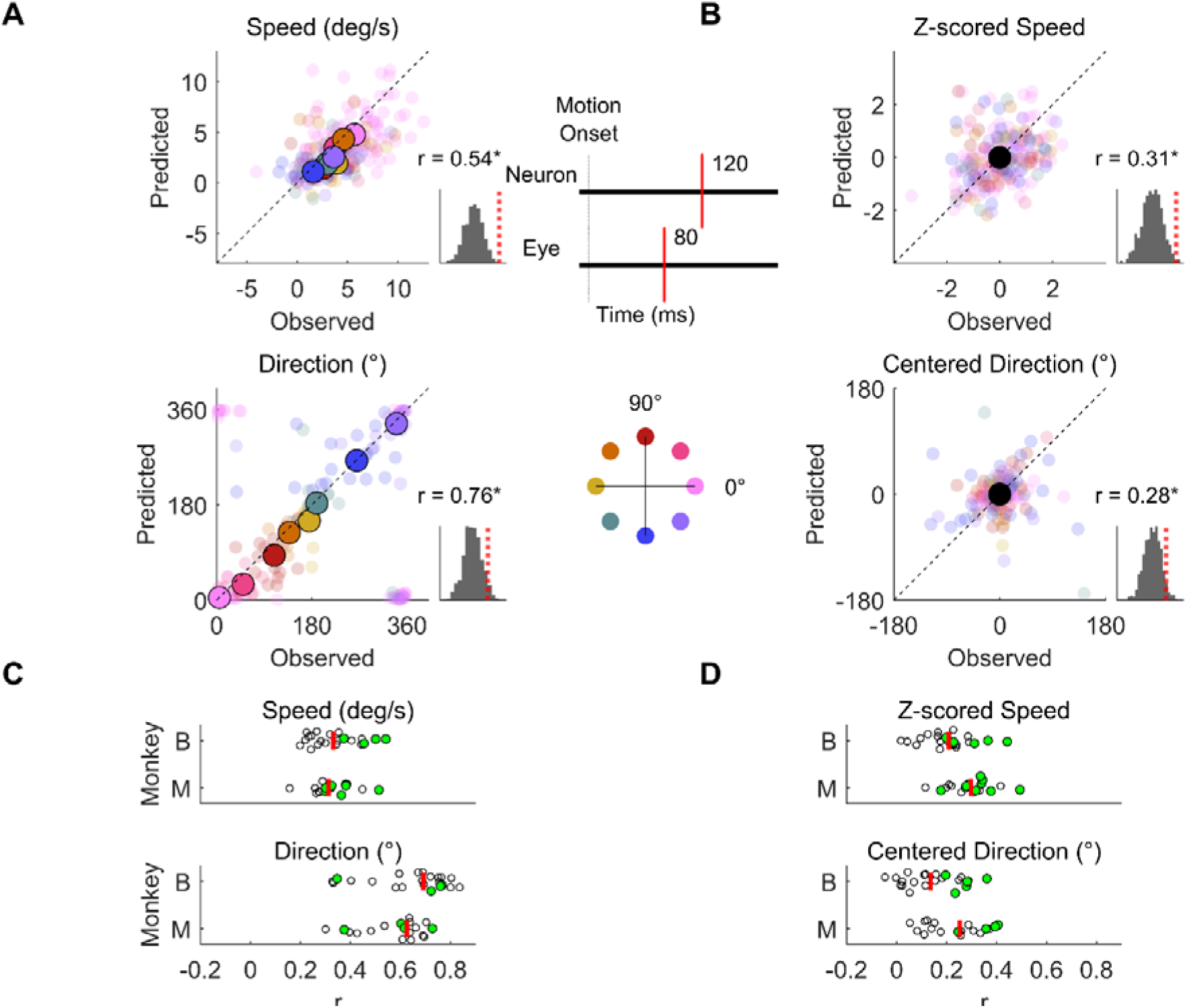
Closed-loop models examining the relationship between early eye movement and later neural activities. (A) Scatter plots illustrating the relationship between observed and PLS reconstructed eye speed (upper panel) and direction (lower panel) at 80 ms, based on neural activity recorded at 120 ms in an example session. Each small dot represents an individual trial, with colors indicating different stimulus directions. Larger dots denote the mean values for each condition. The inset displays the distribution of correlation coefficients obtained from shuffled models, with the red dotted line indicating the experimental values. (B) Scatter plots of predicted and observed eye speed and direction after centering values within each stimulus condition to remove stimulus direction dependent effects. (C) Distribution of correlations in speed and direction in all recording sessions in two animals. Each dot represents one session (n_B_ = 22; n_M_ = 18). Green dots indicate significant correlation in that session. The red bar shows the median across sessions. (D) Similar to (C), but with z-scored speed and centered directions.

Across sessions, we found good performance in the closed-loop models for eye speeds and directions (Figure 2C, D). The mean model performances were significantly above zero in all measures (Monkey B: t = 5.84-14.37, df = 21, all adjusted p < 0.01; Monkey M: t = 8.24-15.75, df = 17, all adjusted p < 0.01; corrected for multiple comparison). For eye speed, we found 18% and 44% of sessions showed significant correlation in Monkey B and M respectively. For direction, we found 18% and 22% significant sessions. After removing stimulus direction dependent effects, we found 23% and 50% significant sessions in z-score speed, as well as 23% and 22% significant sessions in centered direction. In subsequent sections, we will focus the analyses on z-scored speed and centered directions as they are better measures of the trial-by-trial variance in eye movements with the stimulus dependent effect eliminated.

Although model performance was consistently high, it varied considerably across recording sessions. What properties of a recorded neural population determine how well a model performs for that session? To examine this, we ran ANCOVA models with neuronal properties (number of neurons, mean tuning strength or mean circular variance of direction tuning) and animal as predictors (Supplementary Figure 1). Sessions with more units, stronger average tuning, and more distributed preferences for direction yielded better PLS predictions.

### Temporal evolution of relationship between eye movement and neural activities

In the previous analyses correlating eye movements and neural activity, we picked one time point for eye movements (80 ms), and time points before (40 ms, open-loop period) and after (120 ms, closed-loop period) for neural activities. Assuming sensory or motor processing latencies of ∼40 ms, we anticipate that these exemplar open- and closed-loop periods should be most closely related to the eye movements occurring at 80 ms. Given that eye movements and neural activity can mutually influence each other, we next explored the temporal evolution of PLS model performance by reconstructing eye movement at 80 ms with neural activities from 0 to 160 ms with 5-ms steps (Figure 3). We chose a fixed time point for reconstructing eye velocities because it balances being as late as possible to maximize oculomotor gain (i.e. eye speed) while still clearly representing the motor response to the open loop phase of early neural activity. This early time window also minimises confounds associated with saccades, volitional tracking, or deliberately avoiding tracking.

**Figure 3.**
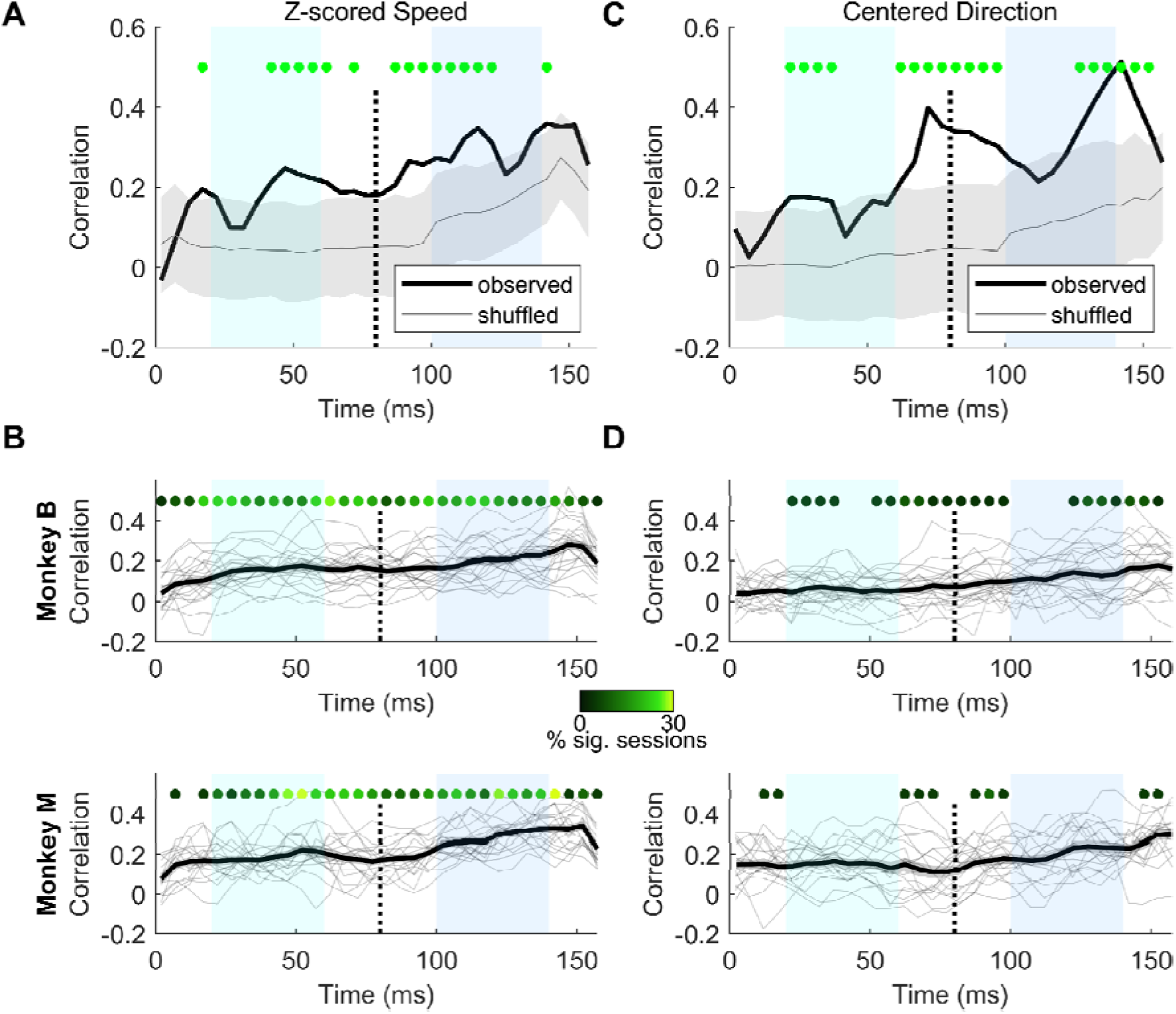
Temporal evolution of model performance of reconstructing eye movement with neural activities at different time points. (A) PLS models were performed with eye movement at 80 ms (dashed line) and neural activities at different time points ranging from 0 to 160 ms with 5 ms time step. Model performance of z-scored speed was plotted against the neural time window used. Light and dark blue regions indicate the exemplar open-loop and closed-loop period for neural activities. The light grey line and grey shaded area shows the mean and 5% - 95% percentile of shuffled data respectively. Significant correlations are indicated with green dots at top. (B) Across session results for both animals. Thin grey lines are the results from individual sessions, and the thick black line is the average across sessions. Graded green dots at the top indicate the percentage of significant sessions at each time point. Absence of dot means no significant results found in all sessions. (C,D) Similar to (A,B) but for centered direction.

We found that model performance for eye speed fluctuates across time in a manner that aligns well with the initial open and late closed loop hypothesis. In the example session (Figure 3A), we found significant models with neural activities at 40-65 ms, which preceded the onset of eye movements. In addition, models with neural activities at 110-125 ms were also significant, reflecting the effect of eye speed on neuronal responses. The pattern was consistent across sessions in both animals (Figure 3B), where we found higher portions of significant models at when using neural activities within the open-loop and closed-loop periods (Monkey B: 18% at 55-60 ms, 23% at 115-120 ms; Monkey M: 22% at 45-50 ms, 17% at 125-130 ms). For centered direction, we found significant models with neural activities at the open-loop period at 20-40 ms, and some significant effects at the end of the closed loop period (125-150 ms) in the example session (Figure 3C). We also found unexpected high model performances with neural activities at 70-100 ms. However, this pattern is not consistent across sessions, with only a limited portion of significant sessions (5-9%) in any time window in both animals (Figure 3D). Overall, the temporal evolution of model performance matched our open-loop and closed-loop hypotheses for eye speed but not for directions.

### Eye and neural time windows for optimal open-loop and closed-loop model performance

Our models focused on exemplar open-loop and closed-loop periods successfully reconstructed eye movements at a predetermined time window at 80 ms. We next explored a broader range of models, allowing offsets of up to 90 ms between the eye and time windows. Correlation values for z-scored eye speed and centered direction were visualized as heatmaps for the same representative sessions from each animal shown previously (Figure 4A, B) and averaged across all sessions for each animal (Figure 4C, D). Significance of correlation values in each individual session was corrected for multiple comparisons with Benjamini-Hochberg procedure.

**Figure 4.**
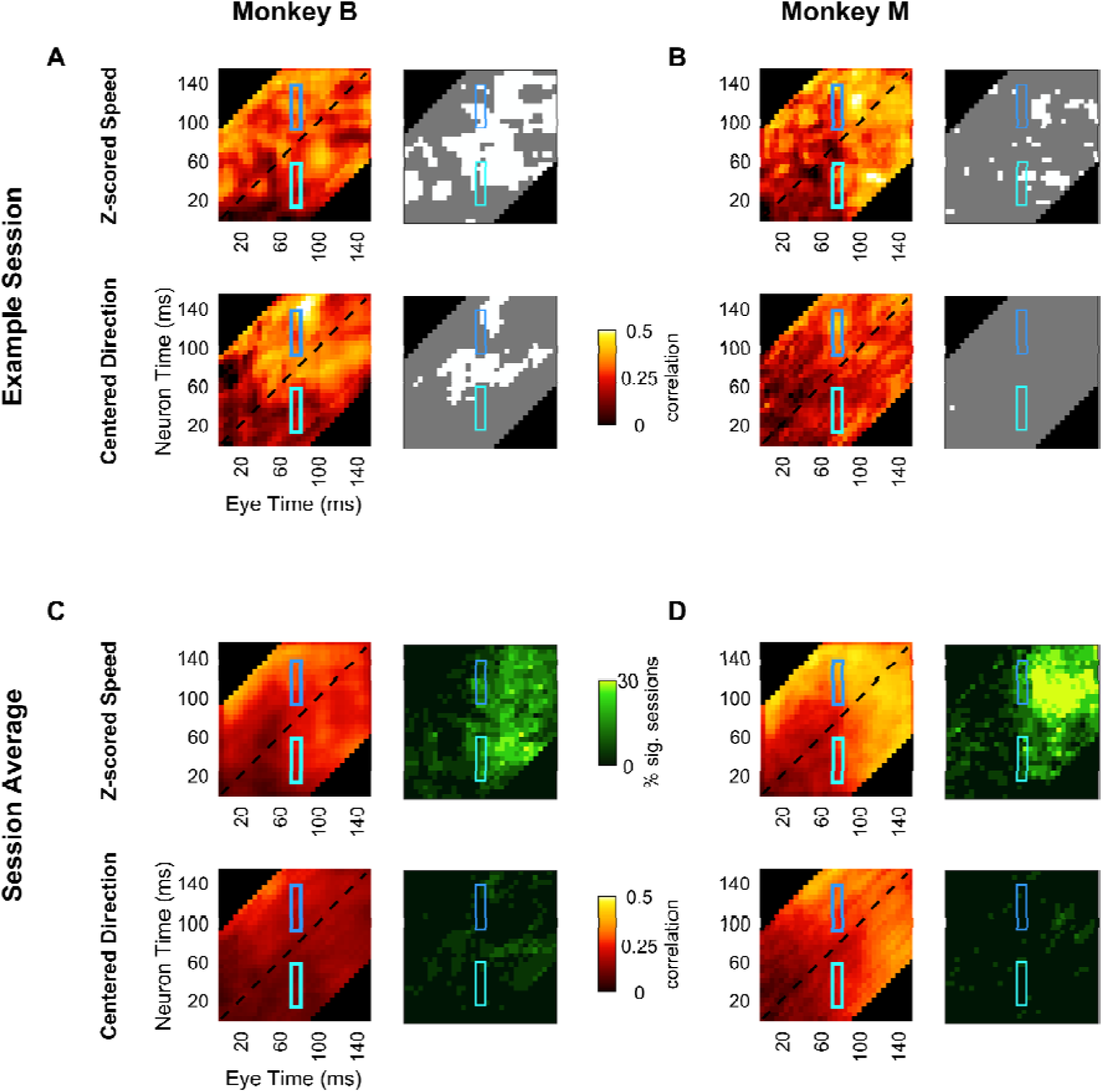
PLS models conducted with a broad range of time points for eye movement and neural activities. (A, B) Heatmaps depicting the correlation between reconstructed and observed z-scored eye speed and centered direction across a range of eye and neural times for an example session from Monkey B (A) and Monkey M (B). Each pixel represents the correlation value for a model using neural activity from a given time point (y-axis) to reconstruct eye movement at another time point (x-axis). The light and dark blue regions highlight the open-loop and closed-loop periods examined in Figure 3. In each panel, grayscale maps on the right show significance, with white regions indicating statistically significant correlations (p < 0.05) and grey regions indicating non-significant correlations. (C, D) Mean heat map for z-scored speed and centered direction across sessions (n_B_ = 22; n_M_ = 18). The maps on the right show the percentage of significant sessions.

The models work best with eye times at ∼100 ms, regardless of the neuronal time window used for reconstruction. In the example session in Monkey B (Figure 4A), we found the peak open-loop model performance for z-scored eye speed at 60/105 ms neural/eye times (r = 0.42, p < 0.01) and peak closed-loop model performance at 145/95 ms(r = 0.4, p < 0.01). For centered direction, best open-loop model happened at 75/105 ms (r = 0.4, p < 0.01) and best closed-loop model was found at 140/ 85 ms (r = 0.54, p < 0.01). For Monkey M (Figure 4B), peak open-loop and closed loop model performance for z-scored speed were found at 45/110 ms (r = 0.55, p < 0.01) and 120/95 ms (r = 0.53, p < 0.01) respectively. No significant results were found in any open-loop or closed-loop models for centered direction. The pattern holds for across session results in z-scored speed, with 27% sessions in Monkey B (Figure 4C) peak at 60/100 ms (mean r = 0.21) for the open-loop models and 18% sessions peak at 120/70 ms (mean r = 0.22) for the closed-loop models. For Monkey M (Figure 4D), open-loop and closed-loop models with most significant sessions were found at 60/110 ms (27%, mean r = 0.27) and 125/90 ms (44%, mean r = 0.34) respectively. For centered direction, limited portions of significant sessions (5-11%) were found in any combination of eye and neural time windows in both animals.

There were high but not significant correlations with eye time 0-60 ms and neural time 90-160 ms (the top left part in the heatmap). A potential cause is the effect of the saccade before stimulus movement, which allows eye speed at time close to 0 to affect later neural activities. However, these correlations also exist in the trial-shuffled distribution as they failed to pass the significance test.

### Latent structures in PLS and PCR are poorly aligned

PLS is a supervised dimensionality reduction technique that identifies latent components that capture the shared variance between the neural and behavioural space. A common alternative to PLS is Principal Component Regression (PCR), which performs regression with principal components defined only on neural activity. While PCR captures variability in the neural space, it’s not clear if this is appropriate for predicting behaviour or is aligned with task demands.

We compared the performance of PLS and PCR to assess whether the latent components explaining both neural activity and behaviour (from PLS) are aligned with the latent components that only explain the space of sensory neuronal responses (from PCR). In our comparison, we focused on the open loop period, where neural activity precedes eye movement by 40 ms. Three key observations can be made in this comparison. First, PLS always outperformed PCR with a matched number of components (t-test, all p<0.05) (Figure 5A-D). This indicates that solely capturing neural variability is not sufficient to predict variations in behaviour. Second, PCR model performance increased very slowly with the addition of more components, such that in all cases, a 5-component PLS model exceeded the performance of a 30-component PCR model. Third, the weak performance of the PCR model is not simply because PCR inadequately captured neural variance. PCR explained similar, but significantly higher, amounts of neural variance than PLS with any given number of components (t-test, all p<0.01; Figure 5E-F).

**Figure 5.**
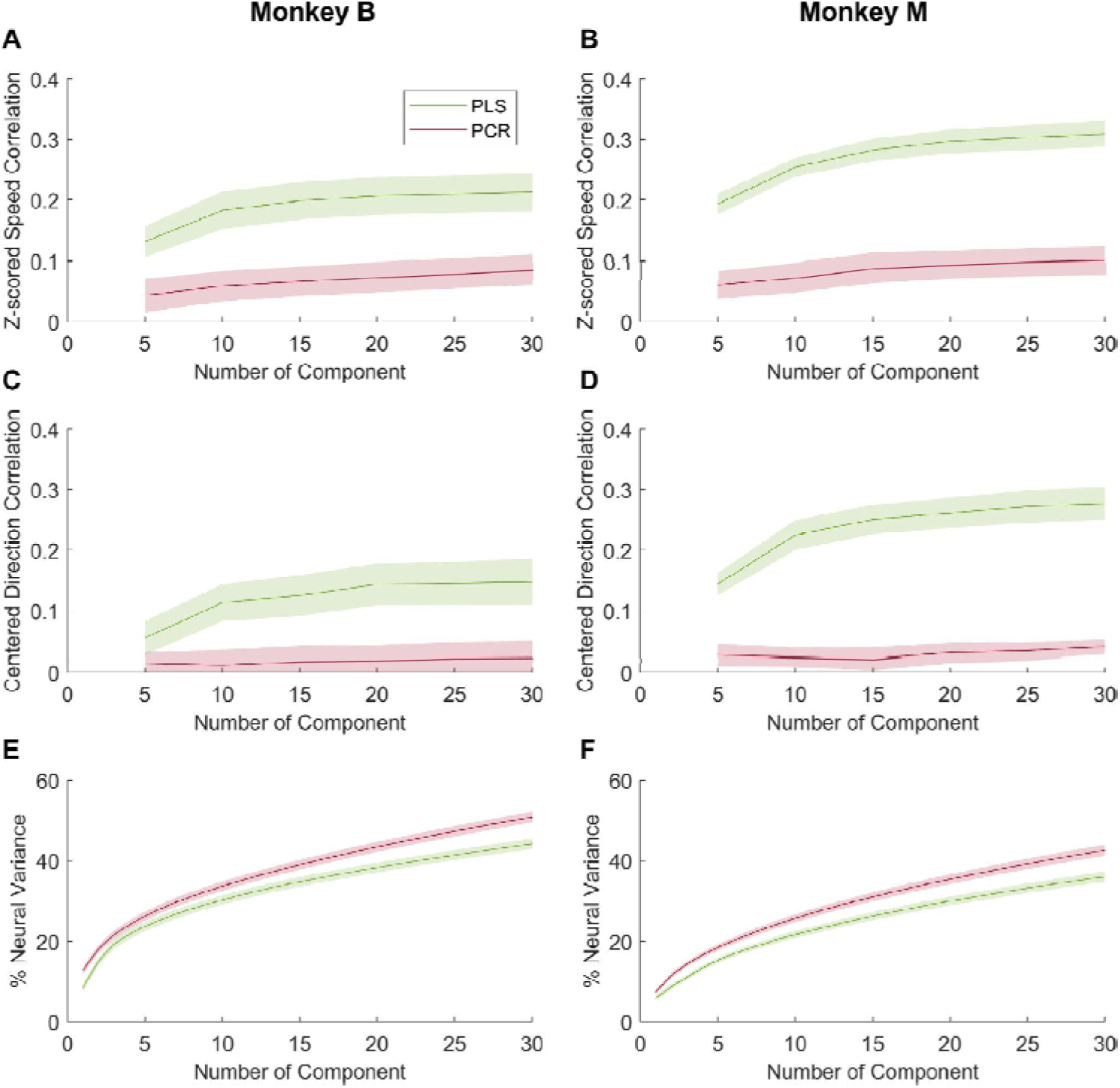
PLS outperformed PCR. (A,B) Model performance of PLS (green) and PCR (red) in z-scored speed across different numbers of latent components. The solid line shows the mean correlation value across all sessions, while the shaded area represents the standard error estimates. (C,D) Similar to A and B, but for centered direction. (E, F) Cumulative percentage of variance explained by PLS and PCR components.

Why does the variance explained by PCR not translate to explaining behavioural variance? To test this, we used Variable Importance in Projection (VIP) scores to characterise how much each neuron contributed to the PLS predictors explaining variance in the behaviour (Supplementary Figure 2). Units with VIP scores higher than 1 are considered to be important predictors in the model (Chong & Jun, 2005). By correlating each unit’s VIP score with its tuning features, we were able to show that units with high firing rates, strong tuning and short latency contribute most to successful PLS reconstructions. These same features should be captured by PCR. Collectively, this illustrates that large amounts of variance in the sensory neural space do not relate to variability in eye movements. Approaches such as PLS are therefore critical for capturing behaviourally-relevant neural variability.

### Vector averaging fails to reconstruct trial-to-trial variations in eye movements

Previous computational models of smooth pursuit proposed that single-trial eye speed can be reconstructed by decoding MT population response with vector averaging (Hohl et al. 2013). Vector averaging combines MT neuron responses by multiplying each neuron’s firing rate by its preferred direction and speed, and then summing these vectors together. To evaluate this decoding method on our dataset, we restricted our analysis to direction-tuned units. We computed a variant of vector averaging called opponent vector average, where preferred directions are decomposed into sine and cosine components when multiplying by firing rates. Therefore, neurons with opposing directional preferences (separated by 180 degrees) would cancel out when both units fire equally. We did not measure the preferred speed of our recorded units. However, we focus our analysis on only the open-loop model (neural time 40 ms, eye time 80 ms), where the visual motion was constant across trials since stimulus speed was the same and the eyes hadn’t started moving. Therefore, trial-by-trial fluctuation in the vector-averaged MT responses should not be severely impacted by preferred speeds.

Our vector averaging models successfully reconstructed uncentered direction but not speed (Supplementary Figure 3), recapitulating the performance of the PLS model seen in Figure 1C,E. Importantly, whereas the PLS model still predicted trial-by-trial variations after centering direction or z-scoring speed, the vector averaging model failed to reconstruct eye speed or direction. In two representative sessions from Monkey B and M (Figure 6A,B), the correlation between model output and observed data were insignificant and negligible (r = 0.04-0.09, p = 0.25-0.58), with the exception of centered direction in Monkey M (r = 0.17, p < 0.01). The mean correlations across sessions were also insignificant and close to zero in all cases (Figure 6C,D, r = −0.02 - 0.01, p = 0.29-0.88). Overall, vector averaging of MT population response did not show shared trial-by-trial variance with eye movement.

**Figure 6.**
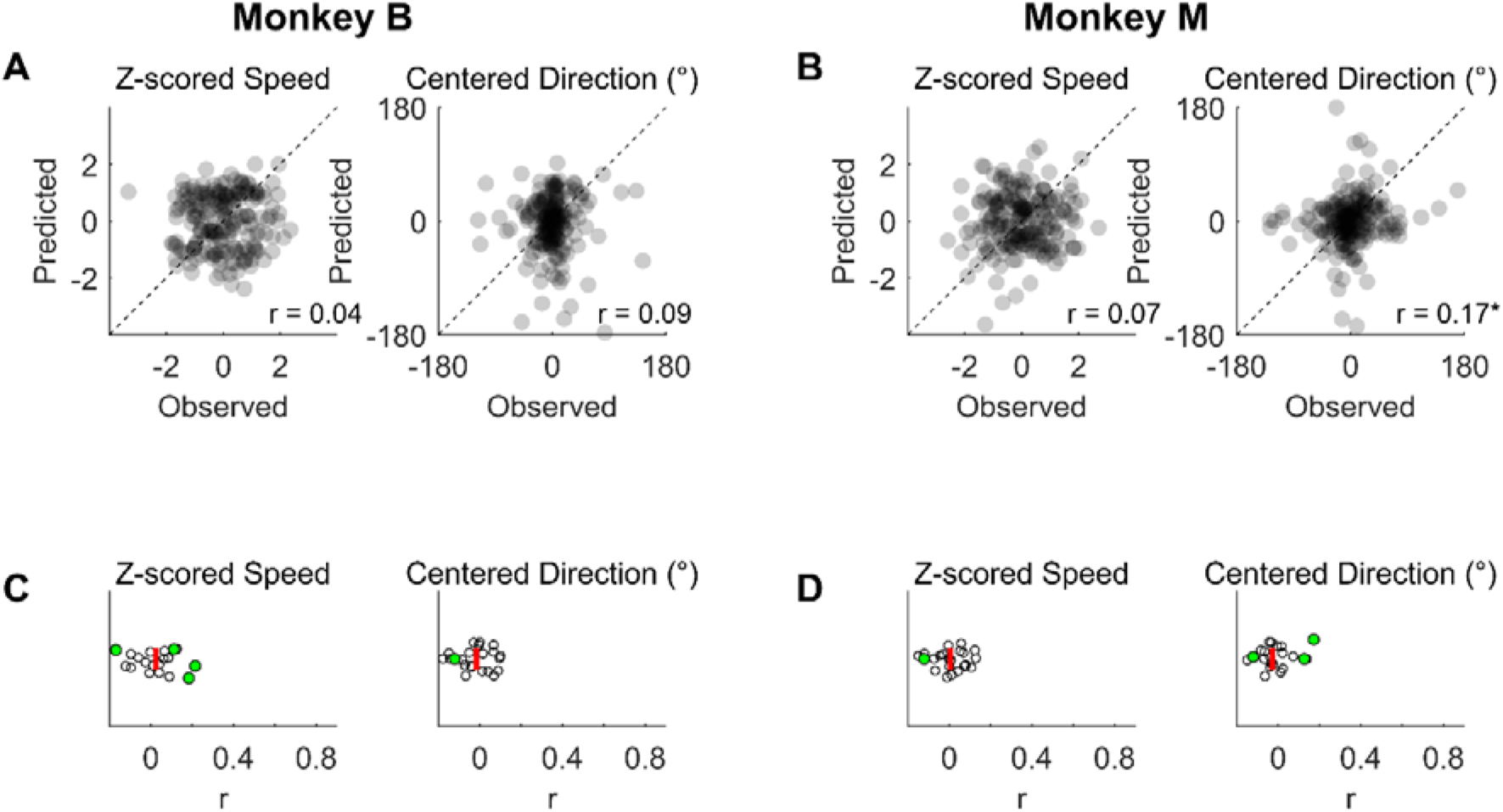
Vector averaging failed to reconstruct eye movements (A) Scatter plots illustrating the relationship between observed and PLS reconstructed eye speed (upper panel) and direction (lower panel) at 80 ms, based on neural activity recorded at 40 ms in an example session of Monkey B. Each small dot represents an individual trial (B) Scatter plot with data from an example session of Monkey M (C) Distribution of correlations in speed and direction in all recording sessions of Monkey B. Each dot represents one session. Green dots indicate significant correlation in that session. The red bar shows the median across sessions. (D) Distribution of correlations across sessions for Monkey M

## Discussion

We characterised the variability shared between neural population activity and reflexive oculomotor responses elicited by moving visual stimuli. We demonstrate that open-loop, stimulus-evoked neural activity could be used to reconstruct eye velocity in subsequent time windows. Given extensive evidence of robust motion-encoding within MT neurons, this was expected because variations in stimulus direction produce large systematic variations in both neural activity and eye velocity. Importantly, our ability to predict eye velocity from neural activity was conserved when we considered responses to repeated presentations of a single motion direction, but only when using a regression approach that jointly considered the neural and motor variability. We used partial least squares regression (PLS), a reduced rank regression approach that finds latent variables based on the space that maximises the covariance between the predictor and observed variables. Therefore, its components account for variability in both neural activity and eye velocity. In contrast, traditional decoding approaches that only apply dimensionality reduction to variability in neural activity, such as principal components regression (PCR) and vector averaging models, were unable to effectively predict eye velocity. This highlights that only some dimensions of neural variability in area MT are aligned with behavioural variability.

Our modeling of the open-loop period of ocular following provides direct support for the hypothesis that the origins of motor variability are variations in the sensory system. Early modeling of the initiation phase of smooth pursuit eye movements suggested that the majority of variance in eye trajectory is attributable to errors in sensory estimates of target velocity (Osborne et al., 2005). This was supported by single-neuron electrophysiological recordings made throughout the sensorimotor hierarchy of macaque monkeys performing pursuit. The average correlation between the eye speed and the firing rate of middle temporal area neurons is 0.1 for neurons with preferred direction aligned with the stimulus direction, and −0.03 for neurons whose nonpreferred direction aligns with the stimulus (Hohl et al. 2013). Later studies showed that neural response latency, but not amplitude, correlates with pursuit latency, and this latency-latency correlation grows throughout the sensorimotor pathway from 0.15 in area MT to 0.37 in floccular Purkinje cells and 0.6 in floccular target neurons and abducens neurons (Lee et al., 2016).

These previous studies suggested that sensory sources (i.e. MT neurons) may account for the majority of variations in pursuit speed and latency. However, they typically recorded only 1 or 2 neurons, forcing them to infer that either the neural population driving pursuit was small, or that neuron-neuron noise correlations in MT were sufficiently large that responses of hundreds of neurons must be decoded downstream to account for motor variations (Huang & Lisberger, 2009; Lee et al., 2016; Shadlen et al., 1996). By recording simultaneously from populations of tens of neurons, we can directly resolve that the latter scenario is more likely. Our finding of a mean correlation of 0.16-0.19 across sessions between predicted and observed eye velocity supports the argument that sensory sources account for some motor variability, but it is much smaller than the levels anticipated if sensory sources account for the bulk of motor variability. Does this amplitude difference reflect differences in task?

The initiation phase of smooth pursuit and ocular following are qualitatively similar, as both involve tracking a moving stimulus. However, there are important differences that may impact the influence of sensory sources. First, ocular following is reflexive, whereas pursuit is a volitional movement. Second, ocular following benefits from post-saccadic enhancement, with increased gain if the stimulus motion triggering tracking occurs shortly after the end of a saccade. However, even with this enhancement, whose underlying mechanisms remain unclear, ocular following tends to have lower open-loop acceleration, lower closed-loop velocity, and therefore lower gain (Mitchell et al., 2015; Pattadkal et al., 2023; Yip et al., 2023). This low gain reduces the signal to noise ratio and therefore reduces the ability to predict eye velocity. However, it cannot account for the differences in decoding performance between PLS, PCR and vector averaging which we also observed.

Previous computational work with simulated neurons suggested that using a vector average approach to decode MT population activity would provide a good estimation of single-trial eye speed (Hohl et al., 2013). In our empirical data, the vector average method performed poorly, potentially because it utilized only the tuning information each neuron conveys and ignored how this information might affect behaviour. We propose an alternative way to approach the question of how to relate variance in the MT population to behavioural variability, by examining the latent neural space that is associated with behaviour. PLS is a supervised method that perfectly suits this framework, because it extracts the shared variance between predictor and response variables. While this means that a given number of latent factors explain less of the neural variance than traditional approaches such as PCR, it performed better in reconstructing eye movements because it explicitly identified a subspace that covaries with both neural activity and behaviour. One limitation of our approach is that we did not completely account for retinal slip variations prior to stimulus motion onset, for example associated with the earlier saccade. This retinal slip may have influenced neural state and the response to the stimulus motion. Regardless, it is still the case that the neural variations we observed only predicted behavioural fluctuations when appropriately decoded.

Choice probabilities have historically been used to quantify how activity of single neurons in MT relates to perceptually-driven behavioural choices (Britten et al., 1996; Price & Born, 2010). A robust finding is that neurons with higher choice probabilities are also more sensitive to the stimuli being judged, initially supporting the idea that there might be alignment between neural representation and behaviour (Pletenev et al., 2026). More recent work suggested that choice probabilities reflect feedback and recurrent processing rather than the impact of feed-forward variability on choices, as choice probabilities grow with stimulus duration and delay, and are impacted by attention (Cumming & Nienborg, 2016; Nienborg & Cumming, 2009). However, the structure of this feedback is not straightforward. When neural populations are considered, population-derived choice probabilities are not well aligned with the stimulus-encoding axis, and population-level choice probabilities can peak after a stimulus ends (Levi et al., 2023; Zhao et al., 2020). The presence of a complex cognitive feedback signal within an ostensibly “sensory” area such as MT is reasonable in the context of a decision making task. Here, however, we study a reflexive oculomotor behaviour and still find that the best representations of sensory-evoked activity are poorly aligned with representations of the neural activity that explain behaviour.

We have previously shown that perceptual judgments and oculomotor responses to the same motion are correlated for smooth pursuit, but weakly or not at all for ocular following (Blum & Price, 2014; Price & Blum, 2014). If a single population of sensory neurons underlies both perception and the eye movements, this suggests that the decoding axes imposed on those sensory signals are poorly aligned. Levi et al. (2023) argued for a form of multiplexed sensory and nonsensory signals in MT that complicates their interaction through simple feedforward and feedback inputs. We further complicate this argument by adding that multiple nonsensory signals and read-out pathways simultaneously exist, as MT must support simultaneous, but distinct, perceptual and oculomotor judgments about the same stimulus. While a significant component of behavioural variability in ocular following can be attributed to variability in sensory representations, it does not reflect the factors of neural activity with the largest variance. A fruitful avenue for future research may therefore be to explore tasks in which sensory signals must be decoded for multiple purposes, such as controlling eye movements and driving perceptual choices. This will allow more direct insight into the multiplexing of sensory, motor and cognitive signals.

## Methods

### Animals and Surgery

Two adult common marmosets (Callithrix jacchus) participated in this study: Monkey B (male, 3 years old) and Monkey M (female, 2 years old). All procedures were approved by the Monash Animal Research Platform Animal Ethics Committee and adhered to the Australian Code of Practice for the Care and Use of Animals for Scientific Purposes. Detailed surgical procedures, including head-post implantation, aseptic techniques, and animal recovery, are described in a previous study (Yip et al., 2023).

To conduct electrophysiological recordings, we performed a craniotomy and implanted a custom-made titanium chamber (internal aperture 12 x 17 mm; height 8.8 mm; weight 6 g) during the same surgery as the head-post implantation. We targeted the center of a 10 x 15 mm craniotomy site at coordinates AP 1 mm, ML 10 mm. After the craniotomy, the recording chamber was glued to the skull with dental acrylic (Ortho-Jet; Lang Dental Mfg. Co.). A thin layer of Kiwk-sil was applied to the craniotomy, and the chamber sealed with a 3D printed resin cap.

### Behavioural Task and Stimulus

During the experiment, the animals sat in a custom-made chair with their heads stabilized by a head-post holder. Eye movements were tracked using the Eyelink 1000 system (SR Research), with binocular tracking for Monkey B (sampling rate 500 Hz) and monocular tracking for Monkey M (left eye, sampling rate 1000 Hz). Eye positions were calibrated with custom scripts at the beginning of each session. To obtain eye velocities from the Eyelink gaze positions, a Savitzky-Golay filter (order 3, length 51 ms) was applied to the raw data.

Stimuli were displayed using MATLAB (MathWorks, Natick, MA) on a 24-inch Viewpixx/3D screen (refresh rate: 100 Hz; resolution: 1920 x 1080) viewed at a distance of 56 cm. Neurostim (https://klabhub.github.io/neurostim/) and the Psychophysics Toolbox extension (Brainard, 1997) were used to generate stimuli and control task sequences.

Within a session, animals completed 100-400 trials of an ocular following task. At the start of each trial, a stationary large-field stimulus (diameter 28 degrees) and a peripheral target (red annulus, diameter 1 degree, with white center, located 5 degrees left from the stimulus center) appeared on the screen. If the animal fixated within 2 degrees of the target for a randomly chosen period of 150-250 ms, the annulus target would disappear and reappear in the stimulus center. The animal was required to make a saccade to the new target location within 800-1000 ms. Upon successful saccade completion, the annulus target would disappear, and after a 50 ms delay the stimulus moved for 300 ms. Afterward, a juice reward was delivered and there was a 1000 ms inter-trial interval. If the animal failed to fixate on the peripheral target within 10 seconds or make a saccade within the time limit, the trial was terminated without a reward.

The stimulus was a “motion cloud” (Leon et al., 2012), a naturalistic grating allowing control of bandwidth of speed, direction, and spatial frequency. Speed (mean 30 deg/s, bandwidth 5 deg/s), spatial frequency (mean 0.15 cpd, bandwidth 0.8 cpd) and direction bandwidth (10°) were fixed across trials and sessions, except for three sessions in Monkey M where the direction bandwidth was reduced to 2°. Note that we use “deg” to refer to spatial parameters and the symbol ° for directions. There were eight possible directions (0-315° in 45° steps), randomized across trials. In 11/22 sessions in Monkey B and 10/18 sessions in Monkey M, there was a 65% probability of direction 0°, while other directions each had a 5% probability (biased-direction task). In other sessions, the 8 directions were equally likely (all-directions task).

### Electrophysiological Recordings

A non-human primate version of the Neuropixels 1.0 probe (Jun et al., 2017) was used for recordings. Reference and ground on the probe were connected and attached to a ground screw in the skull. The 384 channels closest to the tip were recorded on a Windows 10 PC with OpenEphys GUI v0.5.3. The electrode was implanted at the start of each session and extracted at the end of the day. The probe was sterilized with 70% ethanol before recordings. Before each recording session, the craniotomy site was cleaned with peroxide (0.75%), betadine (0.2%), and saline under aseptic conditions. Once the probe had penetrated the dura (confirmed by observing spikes), it was advanced for 1.5-2.8 mm at 4-5 µm/s. The probe was allowed to settle for 20 minutes before starting the ocular following paradigm. After recordings, the probe was extracted and soaked in 1% tergazyme and the craniotomy site cleaning procedure was repeated

Spike sorting was conducted with Kilosort 4 (Pachitariu et al., 2024), and manual curation of spike sorting outputs was done with Phy (https://github.com/cortex-lab/phy). Both well-isolated neurons and multi-unit activities were kept for further analyses. On average, we recorded 350 units (SD = 118) in Monkey B and 504 units (SD = 155) in Monkey M in each session. There were a large portion of multiunit activities (Monkey B: mean = 77.6%, SD = 9.28%; Monkey M: mean = 59.7%, SD = 7.82%) and low firing rate units (<5 spikes/s; Monkey B: mean = 45.74%, SD = 10.14%; Monkey M: mean = 62.47%, SD = 5.21%). The high count of units is probably due to oversplitting, a known issue with modern spike sorting algorithms using template matching methods like Kilosort (Koukuntla et al., 2025). It has less impact in this study because we focus most of our analyses on population analysis, which rely less on perfectly sorted neurons. Moreover, oversplitting should only hurt (e.g. reduce spike count correlations, Cohen & Kohn, 2011) but not artificially boost our model performance.

### Data Inclusion Criteria

Behavioral and neural data quality were checked for each session. To avoid saccade contamination, trials were rejected if an eye speed threshold (30 degrees/s) or position change threshold (3 degrees) were exceeded in the time window 0-160 ms post stimulus motion onset. On average, 84% (SD = 6.48%) trials in Monkey B and 68% (SD = 14%) trials in Monkey M were retained.

For neural data, only units with trial-averaged firing rates (measured at 0-160 ms) higher than 1 spike/s and significant t-test (p<0.05) differences between pre-stimulus (−230 to −100 ms) and post-stimulus (30-160 ms) periods in any stimulus direction were included. An average of 55% (SD = 10%) and 49% (SD = 8%) units per session were kept for Monkey B and Monkey M, respectively.

Sessions were only included if at least 20 direction-selective neurons were recorded with at least an average of 20 trials/direction (all direction task) or 50 trials in direction 0° (biased direction task). We retained a total of 22/28 (Monkey B) and 18/21 (Monkey M) sessions. Average numbers of trials/session were: Monkey B, all-direction task: M = 265, SD = 77; Monkey B, biased-direction task: M = 186, SD = 45; Monkey M, all-direction task: M = 192, SD = 46; Monkey M, biased-direction task: M = 196, SD = 42. Resultant vector length and Rayleigh’s test for nonuniformity were used to determine whether a neuron was direction tuned (circular statistics toolbox, Berens, 2009) .

Significant tuning was defined as having a resultant vector length larger than 0.2 and smaller than 0.8, and a p-value < 0.05 in Rayleigh’s test. Averaged number of tuned neurons were: Monkey B, all-direction task: M = 112, SD = 65 neurons; Monkey B, biased-direction task: mean = 81, SD = 57 neurons; Monkey M, all-direction task: mean = 108, SD = 29 neurons; Monkey M, biased-direction task: mean = 82, SD = 44 neurons.

### Partial Least Square (PLS) Regression

We attempted to reconstruct single trial eye speed and direction with population neural activity at different time windows with PLS regression. Single-trial eye velocities and firing rates were pre-processed to traces with 5 ms resolution. Vertical and horizontal eye velocities at 0-160 ms were discretized into non-overlapping 5 ms bins. Single-trial spike times from each neuron were discretized into 1 ms bins. The spike train was then convolved with a Gaussian kernel with 5 ms standard deviation. Similar to eye velocities, the convolved spike trains were then downsampled to 5 ms resolution.

For each PLS model, population neural activities were defined as predictors and eye velocities as response variables. The segments of eye velocities and firing rates to be selected were determined by a relative offset between the two, ranging from −90 to 90 ms with 5-ms steps. We conducted one PLS model for each offset, resulting in 37 models fitted in each recording session. For example, a 40-ms offset PLS model was performed with single-trial firing rates of all neurons at 0-120 ms and single-trial vertical and horizontal eye velocities at 40-160 ms. Therefore, we had a 3-D matrix of neural activity (Trials x Time x Neurons) and another 3-D matrix for eye velocities (Trials x Time x Horizontal/Vertical). We collapsed trial and time in both neural activities and eye velocities to fit the models. The obtained PLS predictions were then restructured into a 3-D matrix to retain single trial and temporal resolution. We employed MATLAB’s plsregress function, which uses the SIMPLS algorithm (de Jong, 1993). The PLS equation s are:

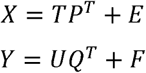

Where X is an n (number of trials) x m (number of units) matrix of predictors and Y is an n x p (number of eye movement measures) matrix of response variables. The matrices T and U, each of dimension n x l (number of latent components), represent the projections of X and Y, respectively – commonly referred to as the X scores (or component/factor matrix) and the Y scores. The loading matrices P and Q correspond to the dimensions m x l and p x l, respectively. Finally E and F denote the residual matrices, which are assumed to follow normal distributions.

To determine the number of components required for each model, an initial PLS with 50 components and 10-fold cross-validation was performed, and the mean squared prediction error (MSE) for each model with XX-YY components was computed. The MSE vs. components line was then fitted with a piecewise regression equation, which has a constant slope before change point c and becomes a flat horizontal line after c. Therefore, the c component would be the first component where increasing the number of components would not further reduce MSE. The mean values of c were 6.61 (SD = 2.29) in Monkey B and 7.23 (SD = 2.8) in Monkey M. PLS was then re-run with c components with 10-fold cross-validation. Predicted values were obtained with the held-out trials.

We converted horizontal and vertical eye velocities in both PLS prediction and observed values into eye speed and direction. We obtained eye directions with the inverse tangent of horizontal and vertical vectors. For eye speed, we noticed that there were biases in the average direction of tracking for different conditions. Therefore, rather than using the raw eye speed (magnitude of the velocity vector) for each trial, we determined the component of eye velocity aligned with the mean tracking response for each condition.

Reconstruction performance was quantified using Pearson correlation between predicted and observed eye speed, and circular-circular correlation (Berens, 2009) between predicted and observed eye directions. To evaluate statistical significance, we performed a permutation-based shuffle test. Specifically, trial labels for eye movement data were randomly shuffled within each stimulus condition 1,000 times, and PLS regression was applied to each shuffled dataset. Predicted eye speed and direction were obtained and correlated with observed data. The resulting distribution of correlation values was then compared to the actual PLS prediction. A prediction was considered statistically significant if it fell above the 95th percentile of the shuffled distribution. Multiple comparisons were corrected using the Benjamini-Hochberg procedure to control for the false discovery rate.

### VIP score analysis

Variable Importance in Projection (VIP) scores were used to evaluate each neuron’s contribution to the PLS models. These represent the normalised variance explained by each PLS component, with a score >1 indicating a highly important variable. First, the PLS weights were normalized:

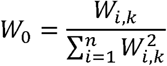

where W is the matrix of obtained PLS weights for predictor i and component k, and n is the total number of predictors. Then, the sum of squares for each component was calculated:

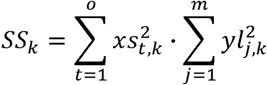

where xs are the scores for component k at the t-th observation, o is the total number of observations, yl are the loadings of component k at the j-th response variable, and m is the total number of response variables. Finally, the VIP score for each predictor was calculated as:

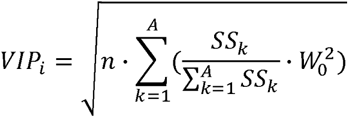

### Comparison of PLS and PCR

We performed Principal Component Regression (PCR) with identical neural data and eye data as the 40 ms offset PLS model. We first performed PCA on the neural data and used PC scores from selected components to predict horizontal and vertical eye velocities with multivariate regression (10-fold cross validation). Eye speeds and directions were then computed as in the PLS analyses. We compared PLS and PCR with 5-30 components, based on the correlation between reconstructed and observed eye speed and direction, and the percentage of neural variance explained.

### Vector averaging

To conduct opponent vector averaging, we first selected only significantly direction tuned units. Then we reconstructed horizontal (V_v_) and vertical (V_v_) eye velocities with the following equations:

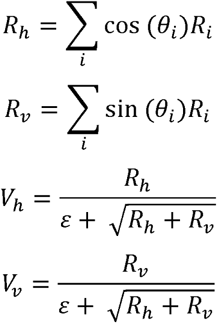

where R_i_ is the single trial firing rate and θ_i_ is the preferred direction of unit i. ε is a constant set at 0.05. Eye speeds and directions were then computed with the predicted vertical and horizontal eye velocities, as in the PLS analyses. The raw eye speeds and directions were further transformed into z-scored speed and centered directions by removing the stimulus condition mean, as done in the PLS analyses. Correlation between reconstructed and observed eye speed and direction were used to evaluate model performance.

## Acknowledgements

This work was supported by Monash Biomedical Imaging Facility and the Monash Animal Research Platform. We are deeply grateful to the dedicated animal care staff for their expertise and commitment and to the two animals whose invaluable contributions made this research possible.

This project was funded by the Australian Research Council (DP200100179; DP2101002107; DP210103865) and by the National Health and Medical Research Council of Australia (APP1185442). M.A.H. was supported by ARC DE180100344; H.M.K.Y. was supported by the Monash Graduate Scholarship.

---

**Supplementary Figure 1.**
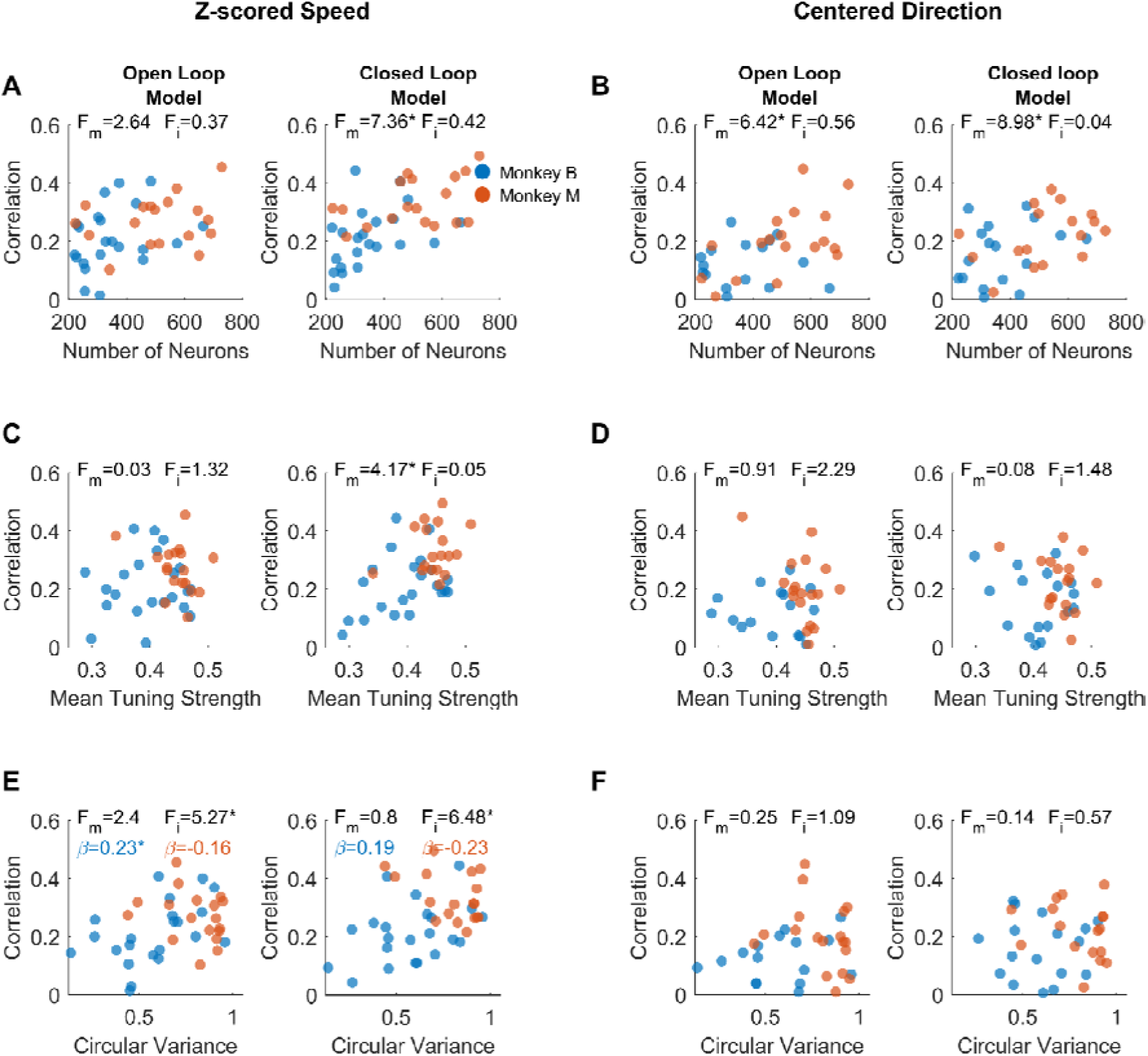
Sessions with more units, stronger tuning and more distributed preferred directions yield better PLS predictions. What properties of a recorded neural population determine how well a model performs for that session? We ran ANCOVA models with neuronal properties (number of neurons, mean tuning strength or mean circular variance of tuning) and animals as predictors. Slopes were allowed to be different between two animals. (A) Scatter plots of number of units vs model performance in z-scored speed in open-loop (left) and closed-loop (right) models. Each dot represents one session, colour coded for two animals. Inset text shows F(1,36) statistics of the main effect of the number of neurons (F_m_) and the interaction term (F_i_) obtained in ANCOVA. Asterisks indicate significance (p<0.05). (B) Similar to (A), but for centered direction (C, D) Scatter plots of mean tuning strength vs PLS predictions. Mean tuning strength is computed as the mean resultant vector length of all direction-tuned neurons. (E, F) Scatter plots of circular variance vs PLS predictions. Circular variance measures the variance of preferred directions in all direction-tuned neurons. Slopes (ß) obtained with simple regression in each animal were denoted when the interaction is significant.

**Supplementary Figure 2.**
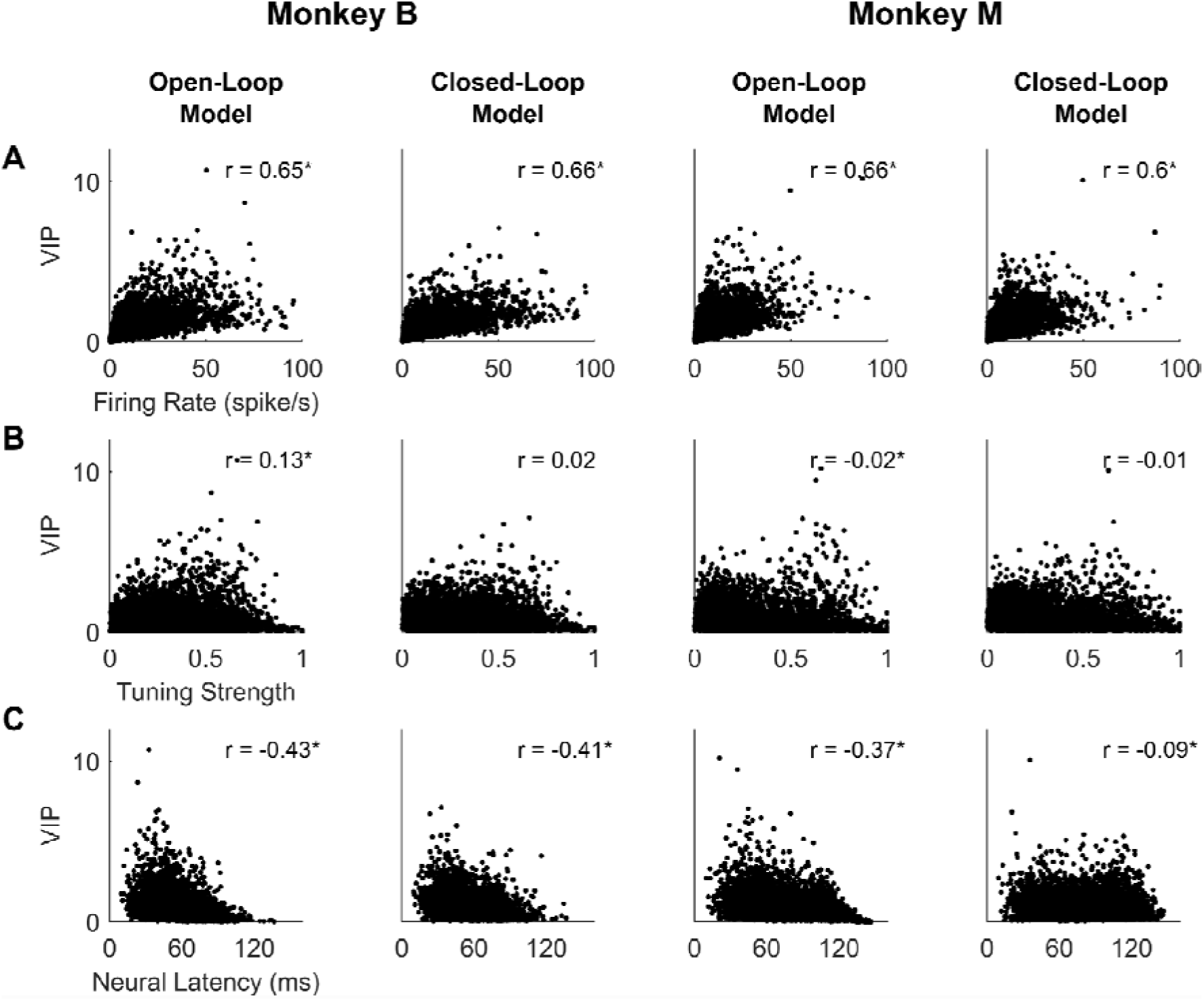
Units with high firing rates, strong tuning and short latency contribute most to PLS reconstructions. Variable Importance in Projection (VIP) scores for neurons quantify the contribution of each predictor to explaining variance in the response variable. Units with VIP scores higher than 1 are considered to be important predictors in the model (Chong & Jun, 2005). VIP scores of individual neurons were plotted against: (A) firing rates 80-120 ms after stimulus onset averaged across all stimulus condition; (B) tuning strength based on the resultant vector length computed from responses to each stimulus direction; and (C) neural latency, determined by averaging single trial latency estimates across all stimulus conditions. We use time points that maximised the number of significantly correlated sessions in Figure 4 (Monkey B: open-loop, 60/100 ms; closed-loop, 120/70 ms; Monkey M: open-loop, 60/110 ms; closed-loop, 125/90 ms). Since VIP evaluates the contribution of individual units, we collapsed data from all sessions in each animal (Monkey B: 7652 units; Monkey M: 8946 units).

**Supplementary Figure 3.**
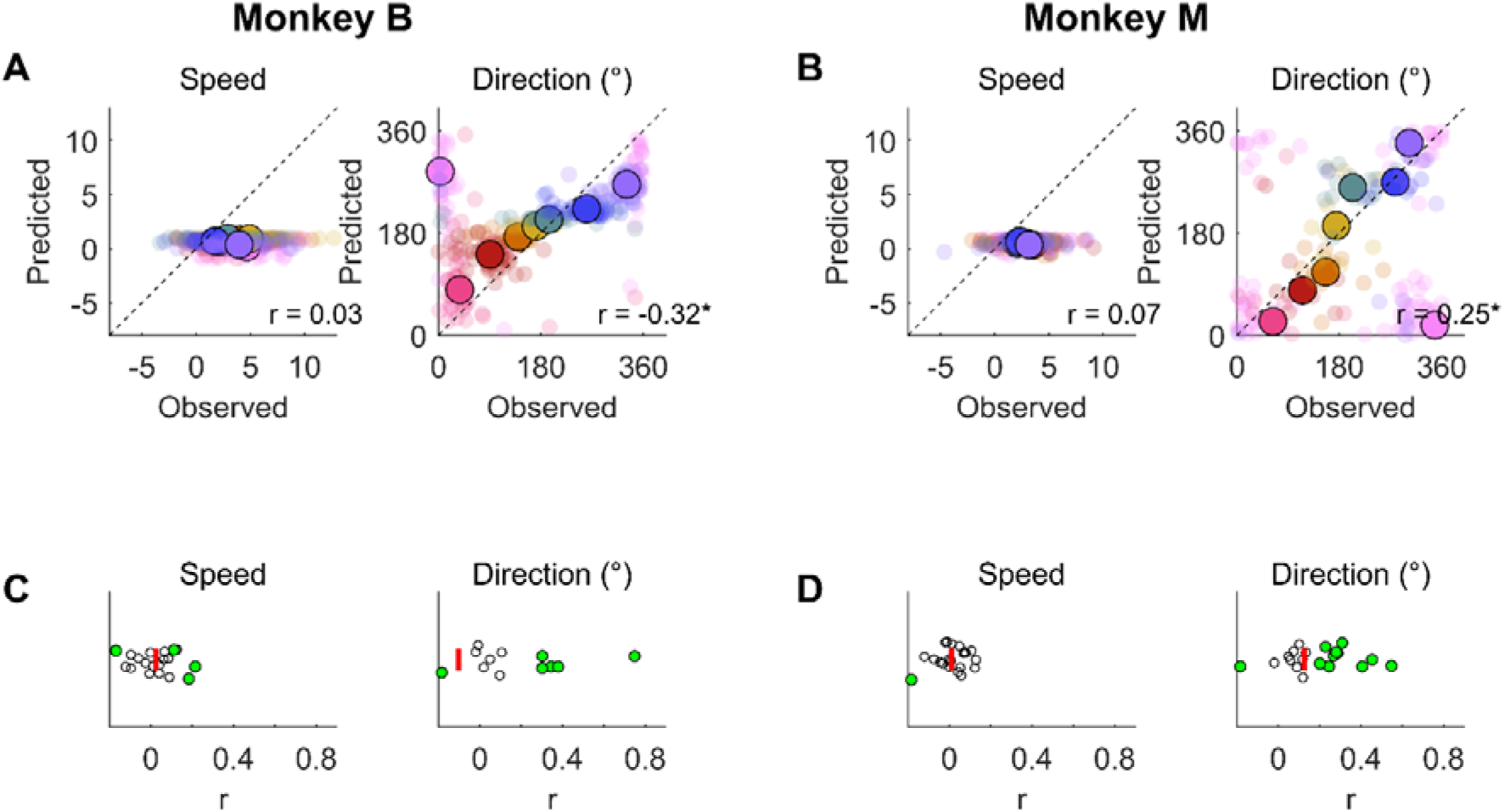
Vector averaging models reconstruct direction but not speed. A vector averaging model was used to reconstruct open-loop eye speed and direction from MT activity before eye movement onset. Same conventions as Figure 1C,E. (A,B) Single-session examples for each animal illustrate good reconstruction performance for direction, but not speed. (C,D) Distribution of correlations in speed and direction in all recording sessions in two animals. Each dot represents one session (nB = 22; nM = 18). Green dots indicate significant correlation in that session. The red bar shows the median across sessions.

